# Integrative Vectors for Regulated Expression of SARS-CoV-2 Proteins Implicated in RNA Metabolism

**DOI:** 10.1101/2020.07.20.211623

**Authors:** Stefan Bresson, Nic Robertson, Emanuela Sani, Tomasz W Turowski, Vadim Shchepachev, Michaela Kompauerova, Christos Spanos, Aleksandra Helwak, David Tollervey

**Author notes:** Equal contributions (Stefan Bresson, Nic Robertson). Correspondence to: Aleksandra Helwak, David Tollervey.

## Abstract

Infection with SARS-CoV-2 is expected to result in substantial reorganization of host cell RNA metabolism. We identified 14 proteins that were predicted to interact with host RNAs or RNA binding proteins, based on published data for SARS-CoV and SARS-CoV-2. Here, we describe a series of affinity-tagged and codon-optimized expression constructs for each of these 14 proteins. Each viral gene was separately tagged at the N-terminus with Flag-His_8_, the C-terminus with His_8_-Flag, or left untagged. The resulting constructs were stably integrated into the HEK293 Flp-In TREx genome. Each viral gene was expressed under the control of an inducible Tet-On promoter, allowing expression levels to be tuned to match physiological conditions during infection. Expression time courses were successfully generated for most of the fusion proteins and quantified by western blot. A few fusion proteins were poorly expressed, whereas others, including Nsp1, Nsp12, and N protein, were toxic unless care was taken to minimize background expression. All plasmids can be obtained from Addgene and cell lines are available. We anticipate that availability of these resources will facilitate a more detailed understanding of coronavirus molecular biology.

## INTRODUCTION

SARS-CoV-2 is a large positive-sense, single-stranded RNA virus that encodes four structural proteins, several accessory proteins, and sixteen nonstructural proteins (nsp1-16) (Figure 1). The latter are mainly engaged in enzymatic activities important for translation and replication of the RNA genome. In addition, nonstructural proteins also target host RNA metabolism in order to manipulate cellular gene expression or facilitate immune evasion (Gordon *et al*, 2020; Thoms *et al*, 2020; Yuen *et al*, 2020). Understanding these pathways in greater detail will be an important step in the development of antiviral therapies.

**Figure 1.**
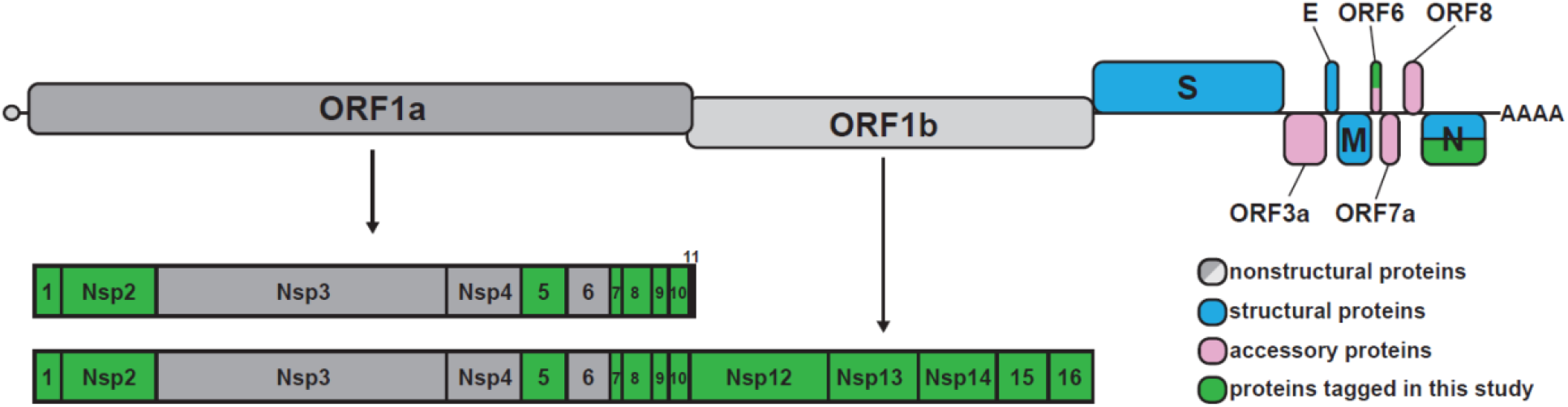
SARS-CoV-2 genome organization. The viral genome consists of a ~30 kb positive-sense transcript that is capped and polyadenylated. Two overlapping open reading frames (ORFs) are translated from the genomic RNA, ORF1a and ORF1ab. Translation of the ORF1b region is mediated by a −1 frameshift allowing readthrough of the stop codon at the end of ORF1a. ORF1a and ORF1ab encode large polyproteins that are cleaved into 16 nonstructural proteins. Structural and accessory proteins (shown in blue and pink, respectively) are separately translated from subgenomic RNAs. ORFs selected for tagging are highlighted in green.

Our lab has previously developed techniques to map the RNA-bound proteome, including <u>c</u>rosslinking and <u>a</u>nalysis of <u>c</u>DNA (CRAC), to identify binding sites for proteins on RNA, and total <u>R</u>NA-<u>a</u>ssociated <u>p</u>roteome <u>p</u>urification (TRAPP) to identify and quantify the RNA-bound proteome (Granneman *et al*, 2009; Shchepachev *et al*, 2019). Among other things, TRAPP has given insights into the mechanism of stress-induced translation shutdown in yeast (Bresson *et al*, 2020). In the context of SARS-CoV-2, CRAC is expected to identify host RNAs that are targeted by viral factors, while TRAPP will reveal how host cell RNA-protein interactions are globally remodeled in response to specific viral proteins.

To facilitate application of these techniques to SARS-CoV-2, we generated a series of synthetic, codon-optimized constructs for 14 different viral proteins that are expected to interact with RNA or RNA binding proteins. To remove the need for error-prone PCR steps, we devised a cloning scheme in which a single synthetic construct could be used to generate untagged, N-, or C-terminally tagged versions of the protein. Using these vectors, we generated and tested a series of human cell lines with the viral ORFs stably integrated, under the control of an inducible Tet-On promoter. We expect that broad availability of this collection will enable a deeper understanding of the coronavirus life cycle and its impact on host RNA biology.

## RESULTS AND DISCUSSION

### Design and cloning of viral expression constructs

We selected 14 proteins for initial analysis, based on putative roles in viral or host RNA metabolism (Figure 1 and Table 1). Two of the selected proteins, Nsp7 and Nsp8, reportedly form a stable heterodimer *in vivo* (Gao *et al*, 2020; Hillen *et al*, 2020; te Velthuis *et al*, 2012), so we also designed constructs in which Nsp7 and Nsp8 were expressed as a fusion protein, connected by a short, unstructured linker.

**Table 1:** Predicted molecular weights and putative functions of selected ORFs

| ORF | Size (aa) | Putative function | References |
| --- | --- | --- | --- |
| Nsp1 | 180 | Suppresses host translation; binds 40S ribosomal subunit | (Huang *et al.*, 2011; Lokugamage *et al.*, 2012; Tanaka *et al.*, 2012; Thoms *et al.*, 2020) |
| Nsp2 | 638 | unknown function; binds eIF4E | (Gordon *et al.*, 2020) |
| Nsp5 | 306 | CL3 M<sub>pro</sub>; Protease (3C-like) | (Kneller *et al.*, 2020; Muramatsu *et al.*, 2016) |
| Nsp7 | 83 | RNA polymerase cofactor | (Gao *et al.*, 2020; Hillen *et al.*, 2020) |
| Nsp8 | 198 | RNA polymerase cofactor; poly(A) polymerase | (Gao *et al.*, 2020; Hillen *et al.*, 2020; Tvarogova *et al.*, 2019) |
| Nsp9 | 113 | ssRNA binding | (Egloff *et al.*, 2004; Littler *et al.*, 2020; Sutton *et al.*, 2004) |
| Nsp10 | 139 | Cap methylation complex | (Chen *et al.*, 2011; Krafcikova *et al.*, 2020; Viswanathan *et al.*, 2020) |
| Nsp12 | 932 | RNA polymerase catalytic subunit | (Gao *et al.*, 2020; Hillen *et al.*, 2020) |
| Nsp13 | 601 | RNA helicase; cap tri-phosphatase | (Jang *et al.*, 2020) (Tanner *et al.*, 2003) |
| Nsp14 | 527 | 3' exoribonuclease (NTD); RNA cap N7 methyltransferase (CTD) | (Ma *et al.*, 2015) (Eckerle *et al.*, 2010; Ogando *et al.*, 2020) |
| Nsp15 | 346 | Uridine-specific RNA endoribonuclease | (Bhardwaj *et al.*, 2004) |
| Nsp16 | 298 | RNA 2'-O-methyltransferase | (Chen *et al.*, 2011; Krafcikova *et al.*, 2020; Viswanathan *et al.*, 2020) |
| Orf6 | 61 | Blocks nuclear-cytoplasmic transport; significantly diverged between SARS-CoV and SARS-CoV-2 | (Frieman *et al*, 2007; Gussow *et al*, 2020; Li *et al*, 2020; Sims *et al*, 2013) |
| N | 418 | Nucleocapsid phosphoprotein | (Cascarina & Ross, 2020; Chang *et al*, 2014; Cubuk *et al*, 2020) |

For cloning, we selected pcDNA5-FRT/TO (Thermo Fisher) as the backbone vector (Figure 2A). This vector can be used for transient transfection or flippase (Flp) recombinase-mediated integration into the genome of cells with a pre-inserted Flp Recombination Target (FRT). It also carries a hygromycin resistance gene to allow selection for integration (Figure 2). As host cells, we used HEK293 Flp-In T-REx cells (293FiTR), but other cell lines that carry an FRT site could also be used.

**Figure 2.**
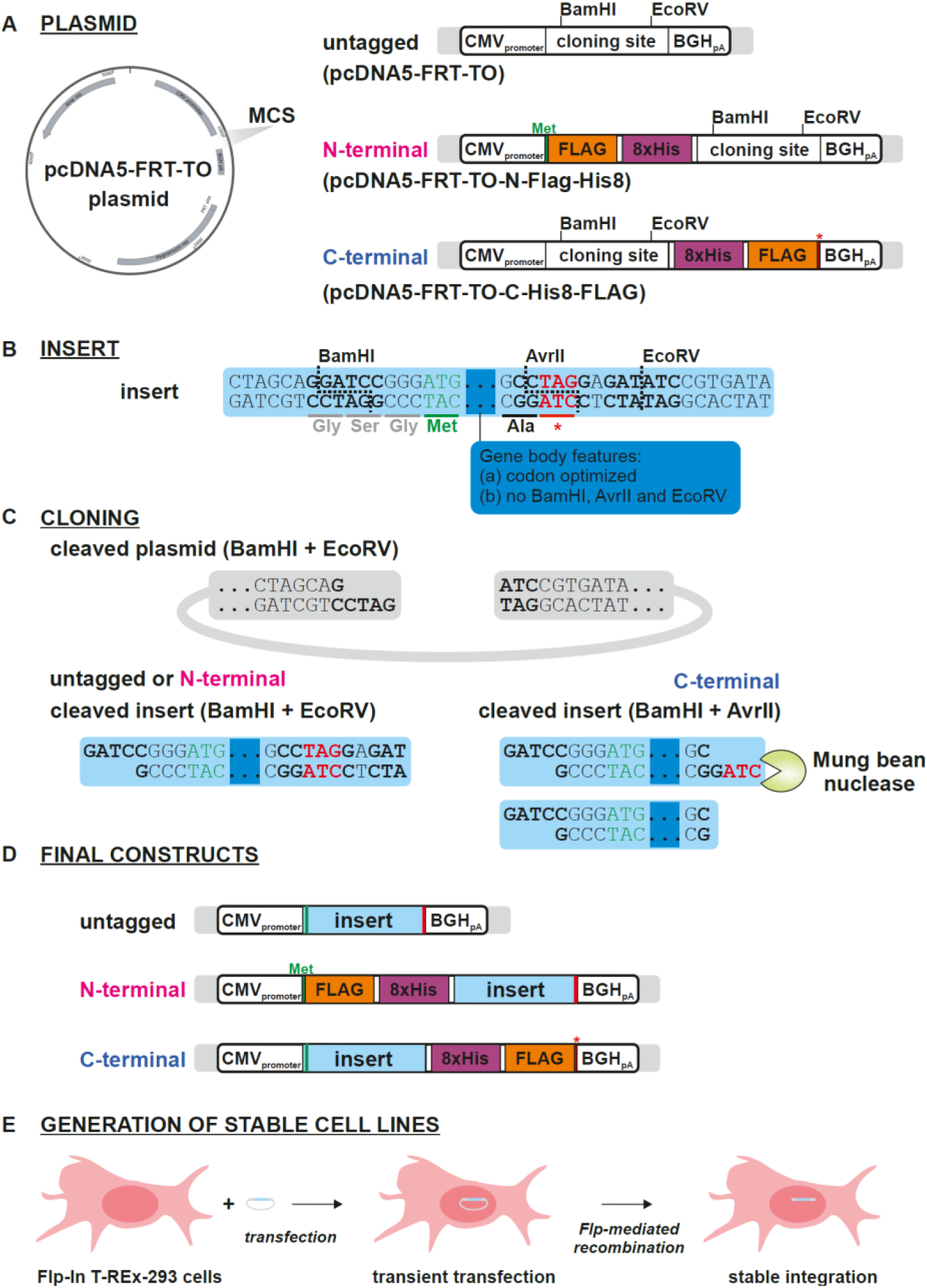
Schematics illustrating the cloning strategy. (A) The three parental vectors used for generating untagged, N-, and C-terminally tagged constructs. (B) Features common to all synthesized insert sequences. Each insert included BamHI and EcoRV sites at either end to facilitate cloning into the three parental vectors. To allow for C-terminal cloning, an AvrII site was inserted such that it overlapped the stop codon (see text for details). In order to accommodate the AvrII site, an alanine residue was added to the end of each expression construct. The viral ORFs were codon-optimized for moderate or high expression, and lacked BamHI, AvrII, and EcoRV sites. (C) For untagged and N-terminal tagging, inserts were digested with BamHI and EcoRV and ligated directly into plasmid precut with the same enzymes. For C-terminal cloning, the inserts were first digested with AvrII, blunted with Mung Bean Nuclease, and then cut with BamHI. The resulting fragment was ligated into plasmid cut with BamHI and EcoRV. (D) Untagged, N-, and C-terminally tagged expression constructs. (E) Strategy for generating stable cell lines (see text for details).

The expression levels of viral proteins vary substantially during infection, so we used constructs that, in addition to stably integrating, were expressed under the control of a tetracycline-regulated human cytomegalovirus (CMV)/ TetO2 promoter, which is induced by the addition of doxycycline to the medium (Figure 2). Viral protein expression can then be titrated by varying either doxycycline concentration or induction time.

Using pcDNA5-FRT/TO as a starting point, we generated two additional parental vectors with pre-inserted tandem-affinity purification tags, either an N-terminal, FH-tag (FLAG-Ala_4_-His_8_) or C-terminal HF-tag (His_8_-Ala_4_-FLAG) (Figure 2A). We have recently shown that these tags work well for tandem affinity purification, including in the denaturing conditions used for CRAC (Bresson *et al*., 2020).

Each synthetic construct was codon-optimized and included a consensus Kozak sequence upstream of the start codon (Figure 2B). The sequences initially used for all ORFs were generated by the algorithms used by Integrated DNA Technologies (IDT; Coralville, Iowa). The prevalence of G-C base pairs, particularly in the third position of codons (GC3), is strongly correlated with increased protein accumulation (Kudla *et al*, 2006; Mordstein *et al*, 2020). To potentially enhance protein synthesis, we ordered alternative ORFs for Nsp8, Nsp13, and N, using the algorithms from GeneArt (Thermo-Fisher Scientific), which have a higher G-C content, particularly in third codon positions.

The insert sequences were designed such that a single synthetic construct could be used to generate an untagged, or N- or C-terminal fusion protein (Figure 2B). BamHI and EcoRV restriction sites were placed on either side of the open reading frame, together with an AvrII site overlapping the stop codon (important for C-terminal cloning, as discussed below).

Generating the untagged and N-terminal tagged constructs was straightforward. The synthetic constructs were cut with BamHI and EcoRV and ligated into pre-cut vector, either pcDNA5-FRT/TO (generating an untagged construct), or pcDNA5-FRT/TO-N-Flag-His8 (generating an N-terminally tagged construct) (Figure 2C). The resulting plasmids were verified by colony PCR and Sanger sequencing.

Generating the C-terminal tagged construct required additional steps to remove the in-frame stop codon upstream of the EcoRV restriction site. Cleavage of the AvrII restriction site, which overlaps the stop codon in the synthetic constructs (Figure 2B), left a 5’ overhang containing the stop codon. Subsequently, the overhang (and thus the stop codon) was removed by treatment with Mung Bean Nuclease, an ssDNA endonuclease (Figure 2C). The resulting fragment possessed a blunt 3’ end, compatible with the EcoRV restriction site in the target vector. Subsequently, the 5’ end of the insert was prepared by digestion with BamHI, and the resulting fragment was ligated into pcDNA5-FRT/TO-C-His8-Flag pre-cut with BamHI and EcoRV.

In total, we generated 54 viral protein expression vectors, and 3 additional GFP expression vectors as controls. These constructs, together with the parental tagging vectors, are listed in Table 2.

**Table 2.** List of available plasmids and cell lines.

| protein | Plasmid number | Tag | Stable cell line | Induction |
| --- | --- | --- | --- | --- |
| pcDNA5-FRT-TO |  | untagged |  |  |
| pcDNA5-FRT-TO-N-Flag-His8 | pSB084 | N-terminal FH |  |  |
| pcDNA5-FRT-TO-C-His8-Flag | pSB085 | C-terminal HF |  |  |
| eGFP | 56 | untagged | yes |  |
| eGFP | 55 | N-terminal FH | yes | +++ |
| eGFP | 57 | C-terminal HF | yes | +++ |
| N protein | 10 | untagged | yes |  |
| N protein | 22 | N-terminal FH | yes | ++ |
| N protein (GC3 opt.) | 28 | untagged | no |  |
| N protein (GC3 opt.) | 31 | N-terminal FH | yes | +++ |
| N protein (GC3 opt.) | 54 | C-terminal HF | yes | +++ |
| Nsp1 | 1 | untagged | no |  |
| Nsp1 | 12 | N-terminal FH | no | N/A |
| Nsp1 | 39 | C-terminal HF | no | N/A |
| Nsp2 | 2 | untagged | yes |  |
| Nsp2 | 33 | N-terminal FH | yes | ++ |
| Nsp2 | 40 | C-terminal HF | yes | +++ |
| Nsp5 | 3 | untagged | yes |  |
| Nsp5 | 13 | N-terminal FH | yes | +++ |
| Nsp5 | 41 | C-terminal HF | yes | +++ |
| Nsp7 | 4 | untagged | yes |  |
| Nsp7 | 14 | N-terminal FH | yes | - |
| Nsp7 | 42 | C-terminal HF | yes | + |
| Nsp7-Nsp8 (GC3 opt.) | 26 | untagged | yes |  |
| Nsp-Nsp8 (GC3 opt.) | 29 | N-terminal FH | yes | + |
| Nsp7-Nsp8 (GC3 opt.) | 53 | C-terminal HF | yes | +++ |
| Nsp8 | 5 | untagged | yes |  |
| Nsp8 | 15 | N-terminal FH | yes | + |
| Nsp8 | 43 | C-terminal HF | yes | - |
| Nsp8 (GC3 opt.) | 35 | untagged | yes |  |
| Nsp8 (GC3 opt.) | 37 | N-terminal FH | yes | + |
| Nsp8 (GC3 opt.) | 50 | C-terminal HF | yes | ++ |
| Nsp8-Nsp7 (GC3 opt.) | 27 | untagged | yes |  |
| Nsp8-Nsp7 (GC3 opt.) | 30 | N-terminal FH | yes | ++ |
| Nsp8-Nsp7 (GC3 opt.) | 52 | C-terminal HF | yes | +++ |
| Nsp9 | 6 | untagged | yes |  |
| Nsp9 | 16 | N-terminal FH | yes | - |
| Nsp9 | 44 | C-terminal HF | yes | + |
| Nsp10 | 7 | untagged | yes |  |
| Nsp10 | 17 | N-terminal FH | yes | - |
| Nsp10 | 45 | C-terminal HF | yes | - |
| Nsp12 | 18 | N-terminal FH | yes |  |
| Nsp12 | 25 | untagged | yes | + |
| Nsp12 | 46 | C-terminal HF | yes | + |
| Nsp13 | 32 | untagged | yes |  |
| Nsp13 | 34 | N-terminal FH | yes | + |
| Nsp13 (GC3 opt.) | 36 | untagged | yes |  |
| Nsp13 (GC3 opt.) | 38 | N-terminal FH | yes | + |
| Nsp13 (GC3 opt.) | 51 | C-terminal HF | yes | +++ |
| Nsp14 | 23 | untagged | yes |  |
| Nsp14 | 19 | N-terminal FH | yes | + |
| Nsp14 | 47 | C-terminal HF | yes | + |
| Nsp15 | 8 | untagged | yes |  |
| Nsp15 | 20 | N-terminal FH | yes | ++ |
| Nsp15 | 48 | C-terminal HF | yes | ++ |
| Nsp16 | 9 | untagged | yes |  |
| Nsp16 | 21 | N-terminal FH | yes | ++ |
| Nsp16 | 49 | C-terminal HF | yes | +++ |
| Orf6 | 11 | untagged | yes |  |
| Orf6 | 24 | N-terminal FH | yes | - |

### Expression of fusion proteins

Each construct was introduced into Flp-In T-REx cells (Thermo Fisher) by transfection, followed by hygromycin treatment for 10-16 days to select for chromosomal integration (Figure 2E). In the initial experiments, all constructs except the Nsp1 series, and the GC3 (high-expression) optimized versions of untagged N and FH-N protein yielded stable hygromycin-resistant cells (Table 2, column 4). Resistant clones were obtained for FH-N by co-transfecting this construct with a plasmid encoding a tetracycline repressor protein (pcDNA6/TR), to suppress possible high levels of FH-N expression at initial transfection.

For expression analyses, doxycycline was added to the cell culture medium to a final concentration of 1 μg ml^−1^, and timepoints were collected for analysis by western blot. A prominent cross-reacting band visible in all samples was used as a loading control for most analyses. The exceptions were Nsp15 and Nsp12, for which anti-M2-Flag and GAPDH were used, as they have migration positions close to the cross-reacting band. Representative data are shown in Figure 3B-C; all other western scans are shown in Figure S1. Quantitation is shown in Figure S2, and associated raw data are presented in Table S4. Clear protein expression was observed for 29 of the 34 tagged constructs (Table 2, column 5). In general, C-terminal tagged proteins were more highly expressed than their N-terminally tagged counterparts.

**Figure 3.**
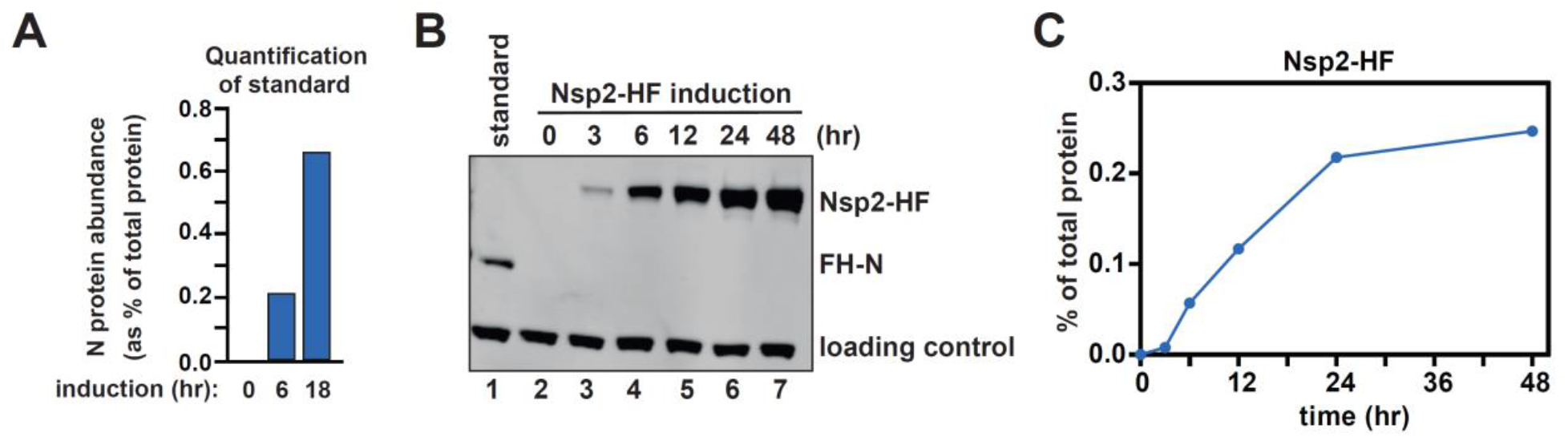
Viral protein induction. (A) Quantification of N protein abundance following a 0, 6, or 18 h induction. Protein levels were assessed by mass spectrometry and label-free quantification, and expressed as a percentage of the total cellular proteome. (B) A representative blot showing induction for C-terminally tagged Nsp2 (lanes 2-7). A prominent cross-reacting band was used as a loading control. To quantify protein abundance, each blot included a normalization standard (lane 1), consisting of lysate from cells expressing N-terminally tagged N protein for 6 h derived from (A). (C) Quantification of the protein bands in (B). Expression was normalized to both the loading control and the N protein standard.

As described above, we were initially unable to generate stable cell lines for any of the Nsp1 constructs. To confirm that Nsp1 could be expressed in cells, we transiently transfected each construct into 293FiTR cells, and confirmed protein expression by western blot. Only the C-terminally tagged Nsp1 showed robust expression (Figure S3).

To allow absolute quantification of SARS-CoV-2 protein expression, we used the N-terminally tagged N protein (integrated from plasmid 22; moderate expression) as a reference standard. Cells containing integrated FH-N were treated with doxycycline for 0, 6, and 18 hours. Protein was extracted, separated by SDS-PAGE and analyzed using mass spectrometry with label-free quantification. To compare proteins, their abundance was expressed as a percentage of the total proteome. This value was calculated using the relative, intensity-based absolute quantification (riBAQ) score for each protein, which represents the iBAQ score for a given protein divided by the iBAQ scores for all proteins. After induction for six hours, N protein comprised 0.022% of the cellular proteome (Figure 3A).

Because all of the tagged proteins possessed an identical FLAG epitope, aliquots of this 6h N protein sample could then be used as a reference standard on all subsequent western blots (e.g. lane 1 in Figure 3B), to allow similar abundance estimates for the other viral proteins (Figure S2 and Table S3). This approach allows viral protein levels to be carefully titrated, so that protein induction approximately matches physiological conditions. This is important because different viral proteins show extreme differences in expression during infection. The nucleocapsid (N) protein can represent 2% of total protein, whereas the non-structural proteins may be 2 to 3 orders of magnitude less abundant (Finkel *et al*, 2020; Grenga *et al*, 2020). Moreover, all proteins will vary in their abundance throughout the course of infection.

### Conclusions

In summary, we report the construction of 57 expression vectors and cell lines. These plasmids can be obtained from Addgene and cell lines are available. We expect that these collections will be a valuable resource for future research into the mechanisms by which coronavirus exploits the genetic machinery of its host to facilitate its own replication.

## MATERIALS AND METHODS

### Construction of expression vectors

All viral genes were cloned into each of three parental vectors: pcDNA5-FRT-TO (to generate an untagged version of the protein), pcDNA5-FRT-TO-N-Flag-His8 (N-terminal tag), and pcDNA5-FRT-TO-C-His8-Flag (C-terminal tag). The N-terminal tag consisted of a single Flag motif, a four-alanine spacer, eight consecutive histidine residues, and a short unstructured linker (DYKDDDDKAAAAHHHHHHHHGSG). The C-terminal tag was essentially the same but in reverse (SGGHHHHHHHHAAAADYKDDDDK).

#### Construction of pcDNA5-FRT-TO-N-Flag-His8

To generate the Flag-His8 tag, we first designed partially complementary DNA oligonucleotides (oSB707 and oSB708) containing the Flag-His sequence. These oligos contained 20 nucleotides of complementarity at their 3’ ends, and an additional 35-36 nucleotides at their 5’ ends. The oligos were annealed in a reaction consisting of 12.5 μM forward oligo and 12.5 μM reverse oligo in a 40 μL reaction volume. The hybridization reaction was initially incubated at 95°C for 6 min, and gradually decreased to 25°C at the rate of 1.33°C/min. Subsequently, the annealed oligos were incubated in the same buffer supplemented with 250 μM dNTPs and 5U Klenow exo- (NEB) in a 50μL reaction at 37°C to fill in the single-stranded regions. After 1 hour, the insert sequence was purified on a silica column (Oligo Clean & Concentrator, Zymo Research).

The insert fragment was then digested with restriction enzymes to facilitate cloning into the acceptor plasmid pcDNA5-FRT-TO. The insert was digested at 37°C for 2 hours in a reaction consisting of 800 ng of DNA, 1X Cutsmart buffer (NEB), 40U HindIII-HF (NEB), and 40U BamHI-HF (NEB). Subsequently, the DNA was purified, as above. In parallel, pcDNA5-FRT-TO was digested under identical conditions but with 2 μg of DNA. The digested DNA was subsequently purified on a silica column (QIAquick PCR Purification Kit; Qiagen). To prevent self-ligation, the vector DNA was phosphatase treated in a reaction consisting of ~2 μg of DNA, 50mM Bis-Tris-Propane-HCl pH 6, 1 mM MgCl_2_, 0.1 mM ZnCl_2_, and 5U of Antarctic phosphatase (NEB) at 37°C for 40 minutes. The DNA was again purified.

Finally, the digested Flag-His insert sequence was ligated into the digested and phosphatase-treated pcDNA5-FRT-TO acceptor plasmid. The ligation reaction consisted of 0.013 pmol vector, 0.042 pmol insert, 50 mM Tris-HCl 7.5, 10 mM MgCl_2_, 1 mM ATP, 10 mM DTT, and 400U of T4 DNA ligase (NEB) in a 20 μL reaction volume. The ligation mix was transformed into homemade DH5α *E. coli*, and plated overnight on LB-Amp. DNA was isolated from several colonies and sequenced to ensure correct insertion of the Flag-His sequence.

#### Construction of pcDNA5-FRT-TO-C-His8-Flag

The pcDNA5-FRT-TO-C-His8-Flag sequence was generated as described above for the N-terminal tagging vector, with the following changes: 1) the oligos used for hybridization were oSB709 and oSB710, and 2) the insert and pcDNA5-FRT-TO were digested with 40U of EcoRV-HF (NEB) and 40U of XhoI (NEB).

#### Cloning of untagged and N-terminally tagged viral genes

pcDNA5-FRT-TO and pcDNA-FRT-TO-N-Flag-His8 were each digested in a 50 μL reaction consisting of 2 μg DNA, 1X Cutsmart buffer, 20U of BamHI-HF, and 20U of EcoRV at 37°C for two hours. In parallel, 2μg of plasmid (KanR) containing the desired viral gene was digested under identical conditions. All three reactions were purified using a PCR cleanup kit. Subsequently, the acceptor plasmids were phosphatase treated as described above and again purified using a PCR cleanup kit.

Digested vector and insert were ligated together in a reaction consisting of 40 ng vector, 120 ng insert, 50 mM Tris-HCl 7.5, 10 mM MgCl_2_, 1 mM ATP, 10 mM DTT, and 400 U of T4 DNA ligase (NEB) in a 10 μL reaction volume. The ligation mix was transformed into homemade DH5α *E. coli*, and plated overnight on LB-Amp. Colony PCR was performed using oAH195-196 to verify the presence of the insert, and positive clones were confirmed by Sanger sequencing.

#### Cloning of C-terminally tagged viral genes

To prepare the backbone, 2 μg of pcDNA5-FRT-TO-C-His8-Flag was digested with BamHI-HF and EcoRV-HF, as above, followed by DNA purification. To prepare viral gene inserts, 1 μg of pUC containing the relevant gene was initially digested with AvrII in the same reaction conditions followed by purification. The 5’ overhang, encoding the stop codon, was then removed by digestion with Mung Bean Nuclease (New England Biolabs): 1 U of Mung Bean Nuclease was added to 1 μg of digested vector in 30 μl of 1x Cutsmart buffer, and incubated for 30 minutes at 30 °C.. After purification, the final insert was produced by digestion with BamHI-HF. The insert was then ligated into the backbone after purification, following procedure described above with a 4:1 molar ratio of insert to backbone.

#### Cloning of eGFP controls

The eGFP insert was amplified using pEGFP-N2 (Clontech) as a template and the DNA oligonucleotides oAH211 and oAH212 (untagged and N-terminal cloning) or oAH211 and oAH213 (C-terminal cloning). Subsequently, the PCR-generated inserts were cloned into pcDNA5-FRT-TO, pcDNA-FRT-TO-N-Flag-His8, and pcDNA5-FRT-TO-C-His8-Flag as described above for the viral constructs.

### Cell culture and transfection

#### Generation of stable cell lines

HEK293 Flip-In TREx cells (Thermo Fisher) were cultured at 37°C with 5% CO_2_ in DMEM (Thermo-Fisher) supplemented with 10% tetracycline-tested FBS (Sigma), 100 μg/mL Zeocin (Thermo-Fisher), and 15 μg/mL Blasticidin S (Sigma). Approximately 1×10_6_ cells were seeded without antibiotics on six-well plates 24 hours prior to transfection. The following day, the viral expression constructs and pOG44 (the FRT recombinase) were co-transfected in a 1:9 ratio (1 μg total) using Lipofectamine 2000 (Thermo-Fisher) according to the manufacturer’s protocol.

The medium was replaced approximately five hours later to remove the transfection reagents. The next day, the cells were split to a 10 cm plate, and after an additional 24 hours, hygromycin B (150 μg/mL) and blasticidin S (15 μg/mL) were added to the medium. Stable integrants were selected over the course of 10-16 days, with medium replacement at regular intervals. Thereafter, stable cell lines were maintained in hygromycin B and blasticidin S.

For the FH-N constructs that initially yielded no colonies, transfection was repeated with the addition of pcDNA6/TR (Invitrogen), encoding the tetracycline repressor protein: total transfected DNA was kept at 1 μg, with pcDNA5/FRT-TO, pcDNA6/TR and pOG44 used at a ratio of 1:4.5:4.5.

#### Transient transfection of Nsp1

Approximately 2×10_5_ cells were seeded without antibiotics on 24-well plates. The following day, 0.2 μg of FH-Nsp1 or Nsp1-HF was transfected into cells using Lipofectamine 2000 (Thermo-Fisher) according to the manufacturer’s protocol.

### Induction tests

For induction tests, 1-2×10_5_ cells were plated into 6 wells each of a 24-well plate. The following day, the cells were induced with 1 μg/mL of doxycycline, and harvested at varying timepoints by resuspending the cells in 50-100 μL of 1X Passive Lysis Buffer (Promega). Inductions were staggered so that all timepoints could be harvested simultaneously. For the transient transfection experiments, cells were induced two hours post-transfection with 1 μg/mL of doxycycline. As with the stable cell lines, inductions were staggered and all timepoints were harvested at once.

For proteins larger than 20 kDa, cell lysates were resolved on 4-12% Bis-Tris gels, run at 180V for 1h in 1X MOPS (ThermoFisher, NP0001). For smaller proteins, lysates were run for 25 min in MES buffer (Thermo Fisher, NP0002). Proteins were wet transferred onto PVDF membrane (Millipore, IPFL00010), blocked with 5% skimmed milk and probed successively with anti-Flag (Agilent, 200474-21) and anti-Rat (Licor, 926-68076). This anti-Flag antibody generate a prominent cross-reacting band at around 50 kDa which was used for loading normalization. For proteins running at the same size of the cross-reacting band (Nsp15 and Nsp12), an anti-M2-FLAG antibody (Sigma-Aldrich, F1804) was used in combination with anti-GAPDH (BioRad, VPA00187) as the loading control. For these, anti-Mouse (Licor, 926-68070) and anti-Rabbit (Licor, 926-32211) were then used. Protein bands were visualized and quantified by scanning in a Licor Odyssey CLX. Normalization was performed using the appropriate loading control and the N protein expression standard which was included on each gel.

### Mass spectrometry

HEK293 cells with stably integrated N-terminal tagged N protein (from plasmid 22) were induced with 1 μg/ml doxycyline for 0, 6, and 18 hours. Cells were harvested in lysis buffer containing 0.1% Rapigest and sonicated (10 cycles of 30 seconds on, 30 seconds off in Bioruper, Diagenode). Approximately 30 μg of protein was resolved on a 4-20% Miniprotean TGX gel, run in Tris-Glycine running buffer at 100 V. Subsequently, the gel was rinsed with water, stained for 1 hour with Imperial Protein Stain (Thermo Scientific), rinsed several times, and destained in water for three hours. Each lane was cut into four fractions and processed using in-gel digestion and the STAGE tip method, as previously described (Bresson *et al*., 2020; Rappsilber *et al*, 2007).

### Data availability

The vectors are available from Addgene: Deposit number 78322. Cell lines are available upon request. The mass spectrometry proteomics data have been deposited to the ProteomeXchange Consortium (http://proteomecentral.proteomexchange.org) via the PRIDE partner repository with the dataset identifier PXD020339 (Perez-Riverol *et al*, 2019).

## ACKNOWLEDGEMENTS

We thank members of the Tollervey lab and Grzegorz Kudla for helpful discussions, This work was supported by Wellcome through a Principle Research Fellowship to D.T. (077248), an ECAT Fellowship to NR (213011/Z/18/Z) and an instrument grant (108504). TWT was supported by the Polish Ministry of Science and Higher Education Mobility Plus program (1069/MOB/2013/0). VS was funded by the Swiss National Science Foundation (P2EZP3_159110). Work in the Wellcome Centre for Cell Biology is supported by a Centre Core grant (203149).

## AUTHOR CONTRIBUTIONS

AH, TT, and SB designed the constructs and cloning strategy. SB, NR, and VS cloned the plasmids. NR, SB, and AH generated stable cell lines. ES, AH, NR, and MK performed western blotting. NR and CS prepared samples for MS analysis. CS and SB performed MS analysis. DT, TT, and AH conceived the project. SB, NR, AH, ES, TT, and DT analyzed data and wrote the manuscript. All authors edited and reviewed the manuscript.

## SUPPLEMENTARY TABLES

**Table S1.** Oligonucleotides used in this work.

**Table S2.** Sequences of fusion protein ORFs.

**Table S3.** Raw data for Figure S2.

**Table S4.** Mass-spectrometry data for N expression.

---

**Figure S1.**
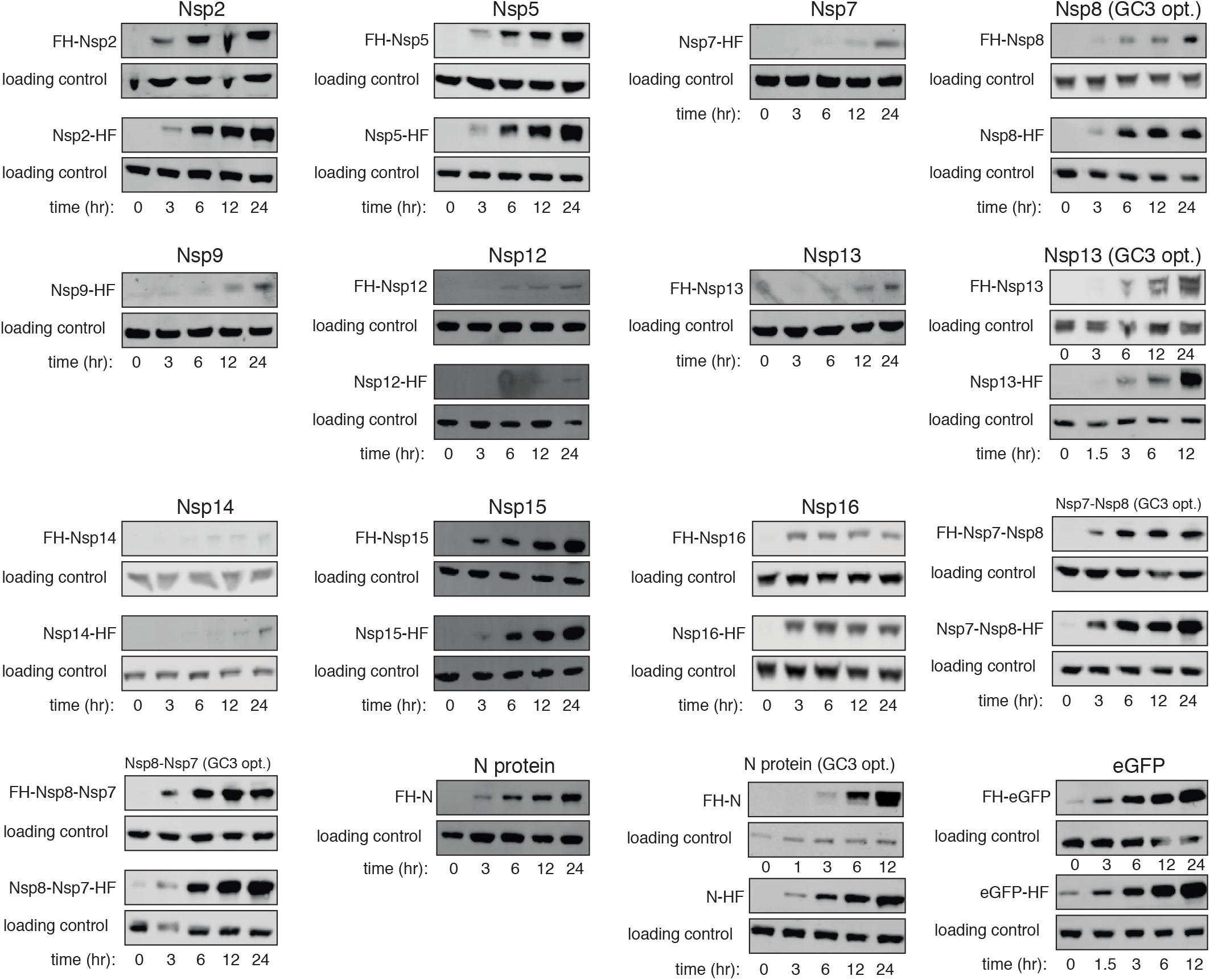
Western analyses of fusion protein expression.

**Figure S2.**
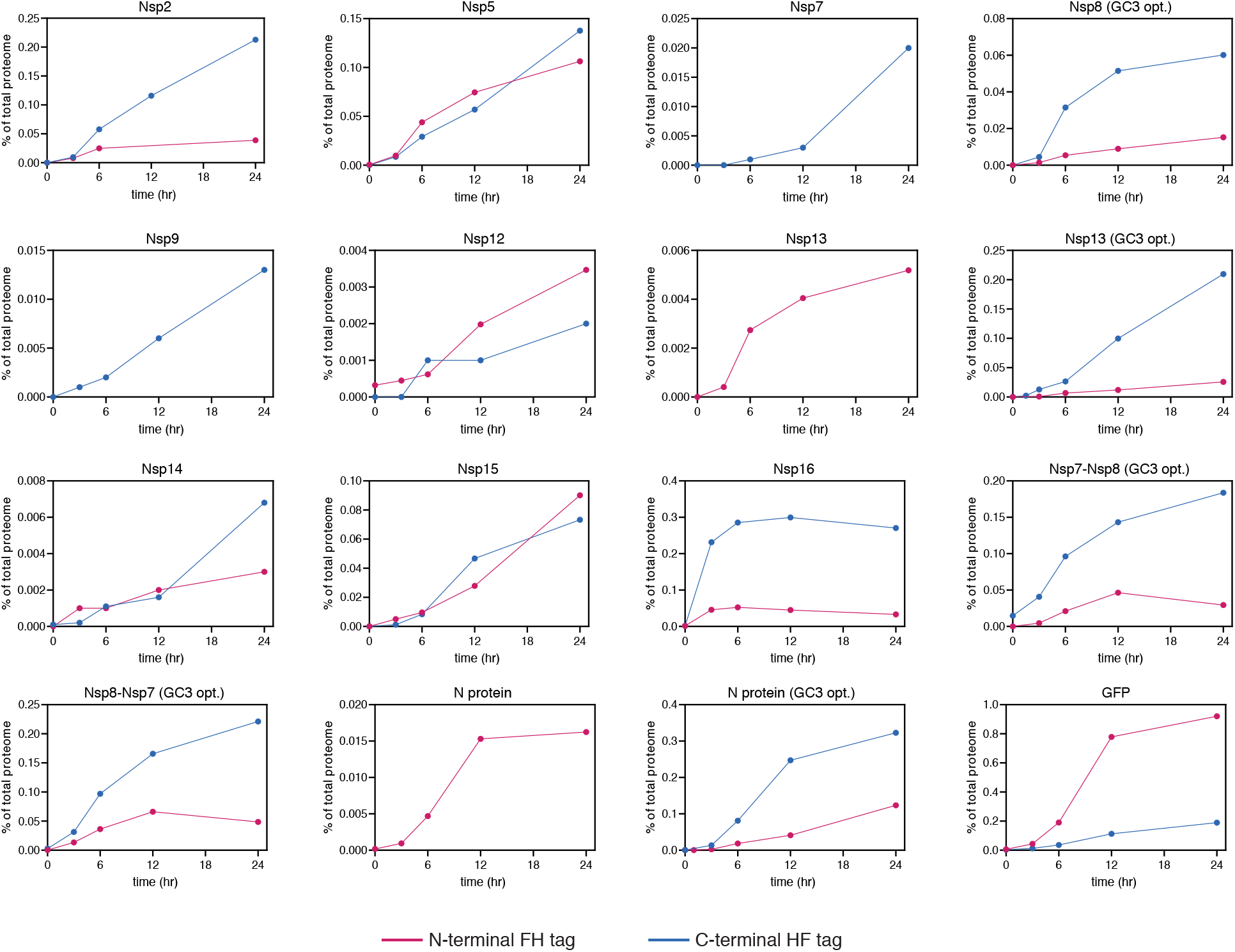
Quantitation of western data from Figure S1.

**Figure S3.**
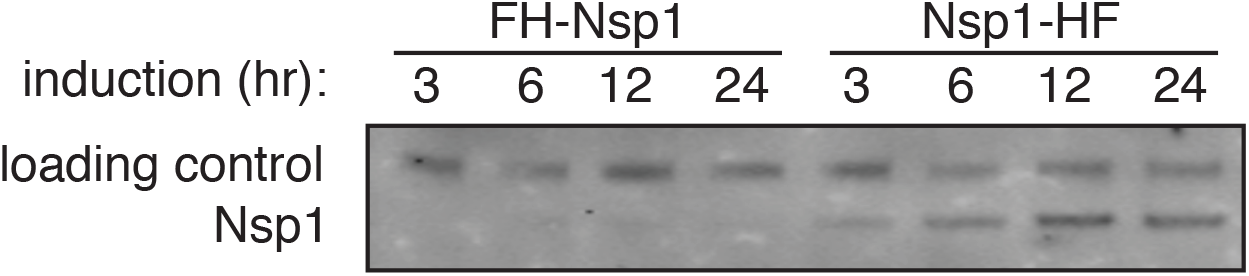
Transient transfection of FH-Nsp1 (derived from plasmid #12) and Nsp1-HF (derived from plasmid #39). Protein expression was assessed following 3 to 24 h induction.

